# Elasmobranchs Exhibit Species-Specific Epidermal Microbiomes Guided by Denticle Topography

**DOI:** 10.1101/2024.04.05.588334

**Authors:** Asha Z. Goodman, Bhavya Papudeshi, Maria Mora, Emma N. Kerr, Melissa Torres, Jennifer Nero Moffatt, Laís F.O. Lima, Ingrid R. Niesman, Isabel Y. Moreno, Michael P. Doane, Elizabeth A. Dinsdale

## Abstract

Elasmobranch epidermal microbiomes are species-specific, yet microbial assembly and retainment drivers are mainly unknown. The contribution of host-derived factors in recruiting an associated microbiome is essential for understanding host-microbe interactions. Here, we focus on the physical aspect of the host skin in structuring microbial communities. Each species of elasmobranch exhibits unique denticle morphology, and we investigate whether microbial communities and functional pathways are correlated with the morphological features or follow the phylogeny of the three species. We extracted and sequenced the DNA from the epidermal microbial communities of three captive shark species: Horn (*Heterodontus francisci*), Leopard (*Triakis semifasciata*), and Swell shark (*Cephaloscyllium ventriosum*) and use electron microscopy to measure the dermal denticle features of each species. Our results outline species-specific microbial communities, as microbiome compositions vary at the phyla level; *C. ventriosum* hosted a higher relative abundance of Pseudomonadota and Bacillota, while *H. francisci* were associated with a higher prevalence of Euryarchaeota and Aquificae, and Bacteroidota and Crenarchaeota were ubiquitous with *T. semifasciata*. Functional pathways performed by each species’ respective microbiome were species-specific metabolic. Microbial genes associated with aminosugars and electron-accepting reactions were correlated with the distance between dermal denticles, whereas desiccation stress genes were only present when the dermal denticle overlapped. Microbial genes associated with Pyrimidines, chemotaxis and virulence followed the phylogeny of the sharks. Several microbial genera display associations that resemble host evolutionary lineage, while others had linear relationships with interdenticle distance. Therefore, denticle morphology was a selective influence for some microbes and functions in the microbiome contributing to the phylosymbiosis.

**Importance:** Microbial communities form species-specific relationships with vertebrate hosts, but the drivers of these relationships remain an outstanding question. We explore the relationship between a physical feature of the host and the microbial community. A distinguishing feature of the subclass Elasmobranchii (sharks, rays, and skates), is the presence of dermal denticles on the skin. These structures protrude through the epidermis providing increased swimming efficiency for the host and an artificial model skin affect microbial recruitment and establishment of cultured microbes but has not been tested on natural microbiomes. Here, we show some naturally occurring microbial genera and functional attributes were correlated with dermal denticle features, suggesting they are one, but not only contributing factor in microbiome structure on benthic sharks.

## Introduction

The coupling of the eukaryotic host and its respective microbial communities ("microbiome") has been reclassified as a meta-organism, representing the mutualistic dependence between a multicellular organism and its respective microbial communities^1^. Factors such as host age^2^, life history ^3^, diet ^4,5^, environment ^6,7^, and body site ^8^ contribute to the composition of microbial communities to varying extents, both internally (e.g., oral, gut, reproductive tracts, etc.) and externally (integumentary system, i.e. skin). As the largest organ of any body, the skin of a host is implicated in numerous facets of host health with the outermost layer of skin, the epidermis, providing a non-invasive avenue to investigate skin health and disease of a host. However, most epidermal microbiome studies focus largely on mammals ^9,10^, particularly humans ^11^. Therefore, characterizing the external microbiomes of non-humans is increasingly necessary to understand how the recruitment and retainment of microbial communities leads to disease.

The epidermal microbiome of different marine vertebrates such as teleost fish, cartilaginous fish (elasmobranchs), and marine mammals contains core microbial species that are conserved throughout their geographic regions and genetically distinction from the surrounding environment^4,12^. Whale sharks (*Rhincodon typus)* from sub-tropical locations around the world share a core microbiome, including the genera *Rheinheimera, Leeuwenhoekiella, Algoriphagus, Sphingoblium, Aeqorivita,* and *Flavobacterium*^5^. Humpback whales (*Megaptera novaeangliae*) found across the northern Pacific, have a distinct set of microbial species including *Psychrobacter*, *Tenacibaculum*, uncultured *Moraxellaceae*, *Flavobacterium*, *Flavobacteriaceae*, and *Gracilibacteri*^13^. Despite the highly diverse surface features of these ocean organisms, each possess unique mechanisms to recruit and maintain respective core microbes. Fish dermis, for instance, feature goblet cells that produce a thick layer of nutrient-rich mucus to cover dermal surfaces^14^, while marine mammals often shed their skin to deter biofilm formation^13^. Elasmobranchs have dermal denticles that protrude through the epidermis to aid in hydrodynamics, while possessing a reduced amount of mucus. These host characteristics exhibit a selective effect with respect to epidermal microbiome structure: the physical characteristic of the shark denticle topography aids in reducing fluid friction and deter biofilm formation while physiological characteristics of the mucus layer provides antimicrobial properties.

The epidermal microbiomes of sharks, which have densely packed denticles, are highly shared across individuals of the same species, while the epidermal microbiomes of stingray, which have sparse dermal denticles and thick mucus, are more variable, suggesting the interaction of dermal denticles and microbiome characteristics ^15,16^. Thus, while microbes pervasively associated with the epidermis of fishes throughout the host’s evolutionary history, describing the specific microbial species present on the skin of marine vertebrates and the host factors influencing microbiome recruitment and retainment is required.

The aforementioned host-microbiota relationships are a product of phylosymbiosis, an outlined trend whereby associated microbial communities are deterministically assembled by host phylogeny across evolutionary time^17^. Examples of these evolutionarily persistent associations include those between coral reef invertebrates^18^, fish^4,19^, and humans ^20^, and their respective microbiomes. A long-standing relationship between a host and microbiome is one that exist between that of sharks and their epidermal microbiome; having an extensive evolutionary history has allowed sharks to adapt alongside a recruited and maintained microbiome and these phylosymbiotic trends occur across shark species including leopard (*Triakis semifasciata*), thresher (*Alopias vulpinus*), blacktip reef (*Carcharhinus melanopterus*), nurse (*Ginglymostoma cirratum*), tiger (*Galeocerdo cuvier*), lemon (*Negaprion brevirostris*), sandbar (*Carcharhinus plumbeus*), Caribbean reef (*Carcharhinus perezii*), and whale sharks (*Rhincodon typus*)^8,19,21–23^. The host-related factors that impact the phylosymbiotic relationship between epidermal microbiomes and shark hosts, remains an outstanding question.

We previously reported the principle that captive, aquatic elasmobranch species maintain a comparable epidermal microbiome to wild counterparts^16^. The nearby proximity of captive elasmobranchs sampled in this study provided an opportunity to characterize the epidermal microbiomes associated with *T. semifasciata*, *H. francisci*, and C*. ventriosum* populations and test whether microbial patterns in benthic shark epidermal microbiomes mirror host phylogeny and correlate with denticle morphology. These sharks were chosen as our model because the hosts are phylogenetically distinct, having an evolutionary distance of approximately 200 million years between *H. francisci* (Heterodontiformes) and both *T. semifasciata* and *C. ventriosu*m (Carcharhiniform)^24^ (Figure 1), are relatively small, can be easily obtained from captive sources, and possess unique dermal denticle topography. We aim to explore whether denticle topography is a feature of the host that is influencing the microbial community and a mechanism underlying emergent patterns of phylosymbiosis. We hypothesize the taxonomic composition of epidermal microbiomes belonging to the benthic elasmobranch species will show correlations both with overall host phylogeny and specific individual microbes. We anticipate the evolutionary lineage of each host species, reflected through alterations in the composition of the microbiome, will also be modulated by the topographical characteristics of the dermal denticles, albeit with weaker covariance.

**Figure 1.**
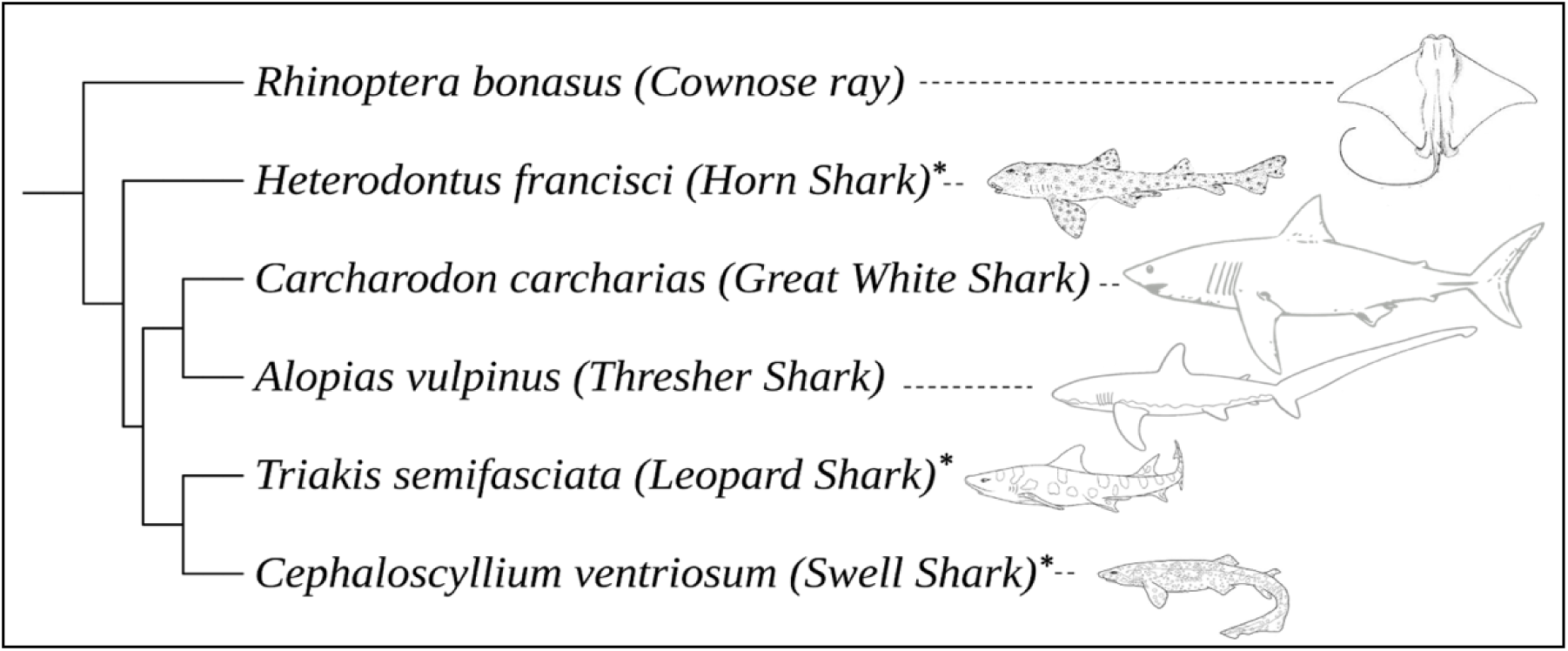
Phylogeny of a subset of elasmobranch species to highlight the relationships of the species investigated in this paper (denoted with an asterisk).

## Results

The epidermal microbiomes belonging to three benthic shark species, *T. semifasciata* (n=11), *H. francisci* (n=10), and *C. ventriosum* (n=10) were sampled in captivity (Table 1) alongside water column samples. Water-associated microbiomes collected from the captive environment were statistically dissimilar to host microbiomes in PERMANOVA main group tests (PERMANOVA: Genus, Pseudo-F _df= 1, 33_ = 3.85, P(perm) < 0.05) and pairwise analyses (*p* < 0.05), did not influence host-associated microbiomes, and are not investigated further.

**Table 1.** Metadata for sampled elasmobranch epidermal microbiomes. Shark species abbreviated as follows: *Heterodontus francisci* (HS, Horn Shark), *Triakis semifasciata* (LS, Leopard Shark), and *Cephaloscyllium ventriosum* (SS, Swell Shark). Sampling environments designated in sample name as Birch Aquarium (BA) or National Oceanic and Atmospheric Administration (NOAA).

| Host Species | Sample Name | Year | Sex | BP | Sequences |
| --- | --- | --- | --- | --- | --- |
| *Heterodontus francisci* | HS BA 1 | 2018 | Female | 105,565,799 | 5,217,911 |
|  | HS BA 2 | 2018 | Male | 82,214,142 | 2,876,236 |
|  | HS BA 3 | 2018 | Female | 74,359,847 | 454,887 |
|  | HS NOAA 1 | 2020 | Female | 79,764,450 | 540,818 |
|  | HS NOAA 2 | 2020 | Female | 34,456,117 | 108,399 |
|  | HS NOAA 3 | 2020 | Female | 165,544,580 | 995,733 |
|  | HS NOAA 4 | 2020 | Female | 143,548,493 | 508,012 |
|  | HS NOAA 5 | 2020 | Female | 133,890,137 | 260,273 |
|  | HS NOAA 6 | 2020 | Female | 112,568,752 | 2,626,021 |
|  | HS NOAA 7 | 2020 | Female | 165,377,448 | 512,408 |
|  | HS NOAA 8 | 2020 | Female | 296,267,509 | 441,426 |
| *Triakis semifasciata* | LS BA 1 | 2018 | Female | 13,254,680 | 1,470,279 |
|  | LS BA 2 | 2018 | Female | 165,572,548 | 295,852 |
|  | LS BA 3 | 2018 | Male | 129,410,990 | 516,511 |
|  | LS BA 4 | 2018 | Female | 140,813,416 | 261,381 |
|  | LS BA 5 | 2019 | Female | 814,847,485 | 3,179,134 |
|  | LS BA 6 | 2019 | Female | 494,586,101 | 246,558 |
|  | LS BA 7 | 2019 | Female | 559,061,883 | 410,944 |
|  | LS BA 8 | 2019 | Female | 451,224,580 | 2,592,630 |
|  | LS BA 9 | 2019 | Female | 877,376,851 | 563,299 |
|  | LS BA 10 | 2019 | Female | 438,923,920 | 429,758 |
| *Cephaloscyllium ventriosum* | SS BA 1 | 2018 | Female | 110,491,568 | 4,549,152 |
|  | SS BA 2 | 2018 | Female | 104,513,010 | 36,040 |
|  | SS BA 3 | 2018 | Female | 146,650,534 | 360,783 |
|  | SS BA 4 | 2018 | Female | 162,351,778 | 490,856 |
|  | SS BA 5 | 2018 | Female | 147,803,072 | 447,888 |
|  | SS BA 6 | 2018 | Male | 170,635,774 | 517,078 |
|  | SS BA 7 | 2018 | Male | 170,164,874 | 320,471 |
|  | SS BA 8 | 2018 | Male | 175,053,487 | 624,057 |
|  | SS BA 9 | 2018 | Male | 502,674,714 | 402,315 |
|  | SS BA 10 | 2018 | Male | 1,097,190,642 | 445,369 |

The total number of bacterial families present for each shark’s microbiome ranged from 164 to 205 individuals associated with *T. semifasciata*, 176 to 205 with *H. francisci*, and 200 to 206 with *C. ventriosum*. To evaluate the overall diversity each of each shark species, the alpha-diversity of the epidermal microbiomes were compared and no differences of microbial community richness (Margalef’s *d*), evenness (Pielou’s *J’*), or overall diversity (Inverse Simpson (1-l)) were observed across shark species (Welch’s t-test; p > 0.05; Table 2) at each taxonomic level (class, family, and genus). The greatest differences observed were measures of richness at genus level, as *C. ventriosum* were recorded to harbor the highest average overall diversity (8.88 ± 1.8 S.D.) followed by *T. semifasciata* (6.36 ± 1.43), and finally *H. francisci* (5.51 ± 1.91).

**Table 2.** Average alpha-diversity metrics of richness, evenness and overall diversity for epidermal microbiome communities associated with *T. semifasciata*, *H. francisci* and *C. ventriosum*.

| Host Species | Margalef's (*d*) Index<br>± S.D. | Pielou's (*J'*) Index<br>± S.D. | Inverse Simpson (1-λ) Index<br>± S.D. |
| --- | --- | --- | --- |
| *T. semifasciata* | 6.36 ± 1.43 | 0.999 ± 1.4E-04 | 0.985 ± 0.002 |
| *H. francisci* | 5.51 ± 1.91 | 0.999 ± 9.5E-05 | 0.979 ± 0.003 |
| *C. ventriosum* | 8.88 ± 1.8 | 0.999 ± 5.9E-05 | 0.992 ± 0.001 |

The benthic sharks microbiome hosted a diversity of bacterial phyla, with the following showing an average relative abundance greater than 1.0%: Euryarchaeota (5.0 ± 1.47 % S.D.; Figure 2), Tenericutes (2.76 ± 0.75 %), Pseudomonadota (2.41 ± 8.82 %), Bacillota (2.37 ± 1.76 %), Actinobactria (2.17 ± 1.26 %), Bacteroidota (1.59 ± 1.97 %), Cyanobacteria (1.58 ± 0.59 %), Aquificae (1.32 ± 0.55 %), Crenarchaeota (1.3 ± 0.9 %), and Spirochaetes (1.23 ± 1.3 %). While each host species harbored the same phyla, the relative abundance varied with *C. ventriosum* retaining the highest relative abundance of Pseudomonadota (39.1 ± 3.25 %) and Bacillota (9.1 ± 3.6 %), while *H. francisci* harbored highest relative abundances of Euyarchaeota (24.5 ± 10.8 %), and Tenericutes (10.8 ± 2.2 %), and *T. semifasciata* highest relative abundance of Actinomycetota (6.32 ± 1.45 %) and Crenarchaeota (5.61 ± 2.67 %).

**Figure 2.**
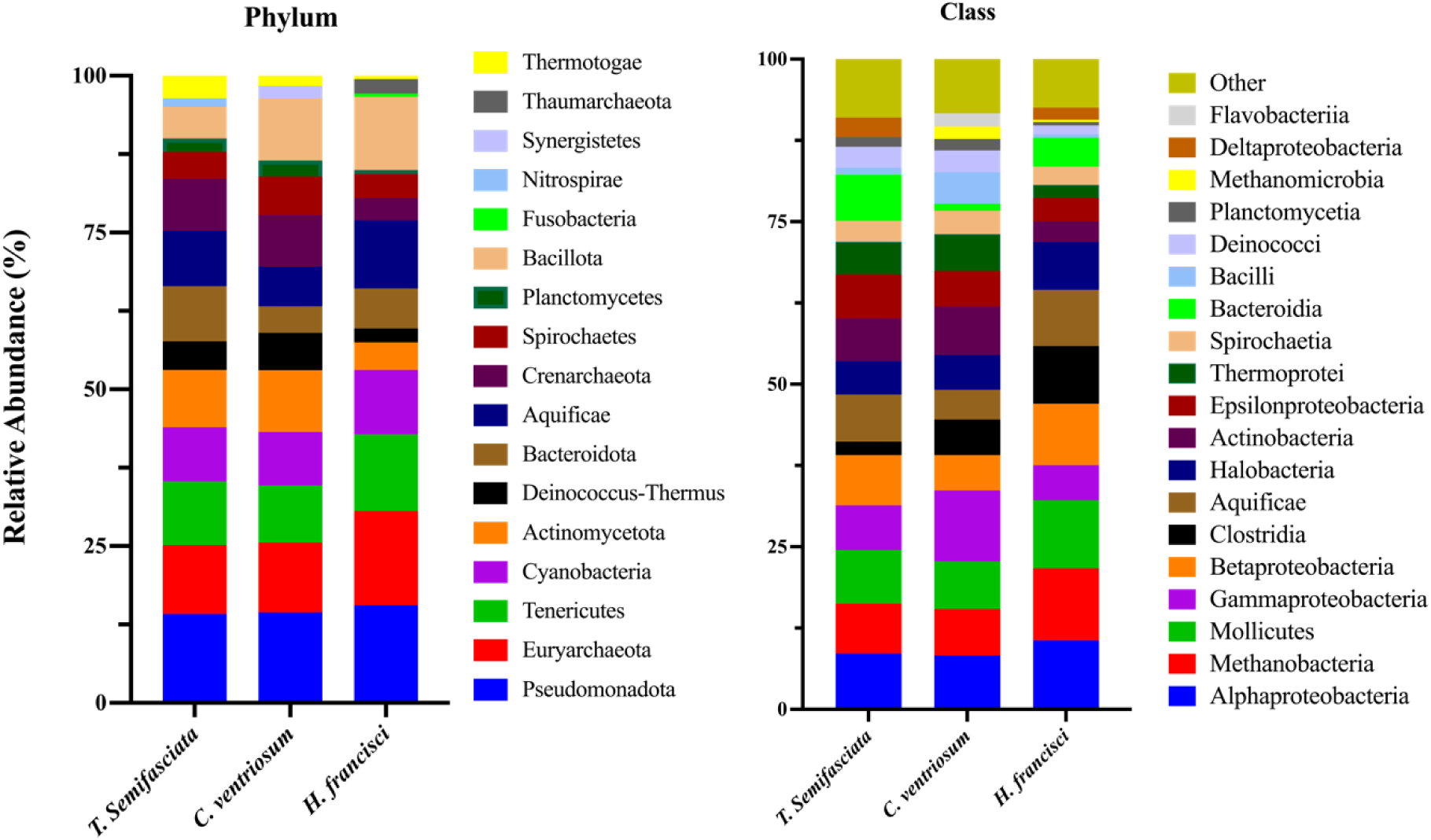
Taxonomic composition of reads from metagenomics sequences from elasmobranch epidermal microbiomes. Left) Relative abundance of microbial phyla identified three benthic shark species. Right) Top 20 class in ascending order based on average by shark species.

The Kruskal-Wallis H test revealed these variations across species to be significant (p < 0.001) between *H. francisci* and both *C. ventriosum* and *T. semifasciata* for the relative abundance of Pseudomonadota and Bacillota. Dissimilarities continued across lower taxonomic levels (all p-values were Bonferroni-corrected). At class taxonomic level, the microbial compositions of microbiomes associated with each benthic shark species exhibited significant differences (p < 0.001) for two taxa: GammaPseudomonadota and Methanobacteria. Significant differences (p < 0.001) between *H. francisci* and *C. ventriosum* included the bacterial classes Actinomycetota, Mollicutes, Aquificae, and BetaPseudomonadota, while differences between *C. ventriosum* and *T. Semifasciata* were observed for BetaPseudomonadota. At genus level, several microbes unambiguously varied between the shark species (Figure 3). For example, the *Bacteriovorax* genus was only present in *T. semifasciata* (2.49 ± 2.15%), while both *Coprococcus* and *Ehrlichia* absent only in *T. semifasciata* epidermal microbiomes. Similarly, *Staphylococcus* was not measured in *H. francisci* epidermal microbiomes.

**Figure 3.**
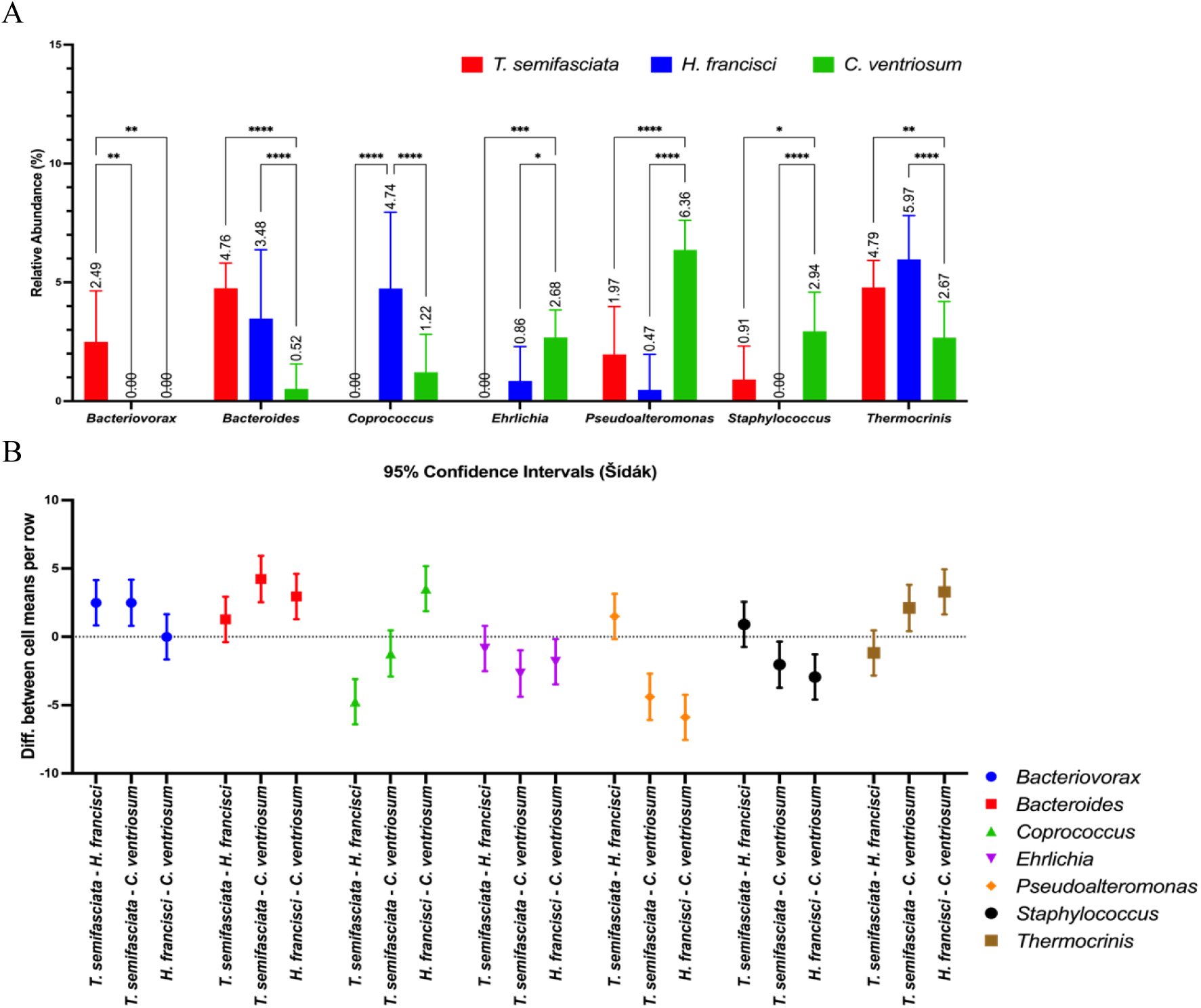
Significant contributors to epidermal microbiomes at A) genus level for associated microbiomes with B) corresponding 95 % confidence intervals.

Pairwise analyses were performed using the Tukey-Kramer post-hoc test at order taxonomic level. The mean abundance order Aquificales on *C. ventriosum* was significantly higher than on both *H. francisci* (p < 0.001) and *T. semifasciata* (p < 0.01). In contrast, for Alteromonadales, the mean abundance in *C. ventriosum* was significantly lower than in both *H. francisci* (p < 0.001) and *T. semifasciata* (p < 0.001). For Campylobacterales, *H. francisci* showed a significantly higher mean abundance compared to both *C. ventriosum* (p < 0.001) and *T. semifasciata* (p < 0.001). Last, for the microbial order Clostridiales, *C. ventriosum* had a higher mean abundance than *H. francisci* (p < 0.001), whereas *H. francisci* showed a lower mean abundance than *T. semifasciata* (*p* < 0.001).

We tested the similarity between and within the groups and identified the key microbes contributing to the dissimilarities between shark species. Comparisons of taxonomic community overlap (SIMPER analysis: 100 - dissimilarity index) within epidermal microbiomes associated with *T. semifasciata*, *H. francisci*, and *C. ventriosum* at family taxonomic level revealed *C. ventriosum* microbiomes to have higher similarity within the sample group (91.9) than those belonging to both *T. semifasciata* (90.4) and *H. francisci* (78.4). The SIMPER analysis for abundance data at genus level reported a dissimilarity coefficient of 62.4 between *C. ventriosum* and *H. francisci*, 58.4 between *T. semifasciata* and *C. ventriosum*, and 53.3 between *T. semifasciata* and *H. francisci*. The most important contributors to the epidermal microbiome dissimilarities between each group varied; microbes belonging to the Bacillota phylum were the top contributors to the differences between *T. semifasciata* and *H. francisci,* Methanobacteria were the most influential contributors to the dissimilarity between *H. francisci* and *C. ventriosum*, and Bacteroidia-encompassing microbes impacted the difference between *T. semifasciata* and *C. ventriosum* (Table 3). In pairwise comparisons between *T. semifasciata* and *H. francisci*, the *Coprococcus* genus was found to contribute the most to the dissimilarity, with a contribution of 4.45 %. This genus was present in T*. semifasciata* but absent in *H. francisci*. Other significant contributors included *Methanothermobacter* (3.69 %) and *Methanobacterium* (3.0 %), with the former being absent in *H. francisci*. When comparing *H. francisci* and *C. ventriosum*, *Methanothermobacter* emerged as the top contributor to the dissimilarity, with a contribution of 6.42 %, as it was more abundant in *C. ventriosum* than in *H. francisci* metagenomes. In the comparison of *T. semifasciata* and *C. ventriosum*, *Methanothermobacter* again contributed the most to the dissimilarity (5.61 %), being more abundant in *T. semifasciata*.

**Table 3.** Pairwise comparison and resulting percent contribution of genera driving differences between epidermal microbiomes associated with each host, determined by SIMPER analysis.

| Host Shark Species | Genus | Contribution (%) | Average Relative Abundance |  |
| --- | --- | --- | --- | --- |
|  |  |  | *T. semifasciata* | *H. francisci* |
| *T. semifasciata*<br>vs<br>*H. francisci* | *Coprococcus* | 4.45 | 0.0 | 4.74 |
|  | *Methanothermobacter* | 3.69 | 0.0 | 3.93 |
|  | *Methanobacterium* | 3.0 | 1.16 | 3.44 |
|  | *Haloquadratum* | 3.01 | 3.27 | 5.32 |
|  | *Jonesia* | 2.91 | 4.6 | 2.01 |
|  | *Caldivirga* | 2.78 | 2.97 | 0.0 |
| *H. francisci*<br>vs<br>*C. ventriosum* |  |  | *C. ventriosum* | *H. francisci* |
|  | *Methanothermobacter* | 6.42 | 9.63 | 4.26 |
|  | *Methanobacterium* | 3.19 | 4.74 | 1.22 |
|  | *Coprococcus* | 3.36 | 5.18 | 1.56 |
|  | *Thermocrinis* | 2.76 | 5.97 | 2.67 |
|  | *Bacteroides* | 2.68 | 3.48 | 0.52 |
|  | *Haloquadratum* | 2.63 | 5.32 | 2.67 |
|  | *Pseudoalteromonas* | 2.62 | 0.23 | 3.5 |
|  | *Tremblaya* | 2.5 | 6.08 | 3.09 |
| *T. semifasciata*<br>vs<br>*C. ventriosum* |  |  | *T. semifasciata* | *C. ventriosum* |
|  | *Methanothermobacter* | 5.61 | 4.61 | 1.3 |
|  | *Bacteroides* | 3.62 | 4.76 | 0.52 |
|  | *Methanobacterium* | 2.35 | 0.0 | 2.86 |
|  | *Pseudoalteromonas* | 2.45 | 3.36 | 1.56 |
|  | *Ehrlichia* | 2.29 | 0.0 | 2.68 |
|  | *Caldivirga* | 2.25 | 2.97 | 1.09 |

We performed PERMANOVA tests on the microbial compositions of each shark to determine if significant differences were present between the epidermal microbiomes associated with each host species. The main PERMANOVA tests identified significant differences in the proportional abundances at family and genus levels (PERMANOVA: family, pseudo-*F* _df= 2, 28_ =7.65, *P*(perm) = 0.001; genus, pseudo-*F* _df= 2, 28_ = 6.79, *P*(perm) = 0.001), while pairwise PERMANOVA tests outlined significant differences (P(perm) ≤ 0.002) between the epidermal microbiome belonging to each species (Table 4) at each taxonomic level (order, family, and genus).

**Table 4.** Summary of pairwise PERMANOVA results for epidermal microbiome compositions between elasmobranch species at order, family, and genus level (999 permutations).

| Host Factors, Taxa | Order Level |  | Family Level |  | Genera Level |  |
| --- | --- | --- | --- | --- | --- | --- |
|  | t(test) | *P*-(perm) | t(test) | *P*-(perm) | t(test) | *P*-(perm) |
| *T. semifasciata* vs *H. francisci* | 2.26 | 0.002 | 2.34 | 0.002 | 2.34 | 0.001 |
| *T. semifasciata* vs *C. ventriosum* | 2.93 | 0.001 | 2.55 | 0.001 | 2.76 | 0.001 |
| *C. ventriosum* vs *H. francisci* | 3.11 | 0.001 | 3.52 | 0.001 | 2.93 | 0.001 |

To clarify the relationships among the epidermal microbiomes of different host species, hierarchical clustering was performed at the genus level for each sample. Using the Bray-Curtis similarity index, the microbial genera within the epidermal microbiomes formed distinct clusters, each corresponding to a unique host species (Figure 4A). An nMDS plot showed further clustering of the metagenomes by host species with a stress value under the acceptable threshold (0.13), suggesting the differences between host species are significant influencers over microbial community structure (Figure 4B). Finally, Spearman’s rank correlation coefficients, based on microbial genera, demonstrated stronger pairwise relationships between *T. semifasciata* and *C. ventriosum* (0.63), than between *H. francisci* (0.56), and between *H. francisci* and *C. ventriosum* (0.61; Figure 4).

**Figure 4.**
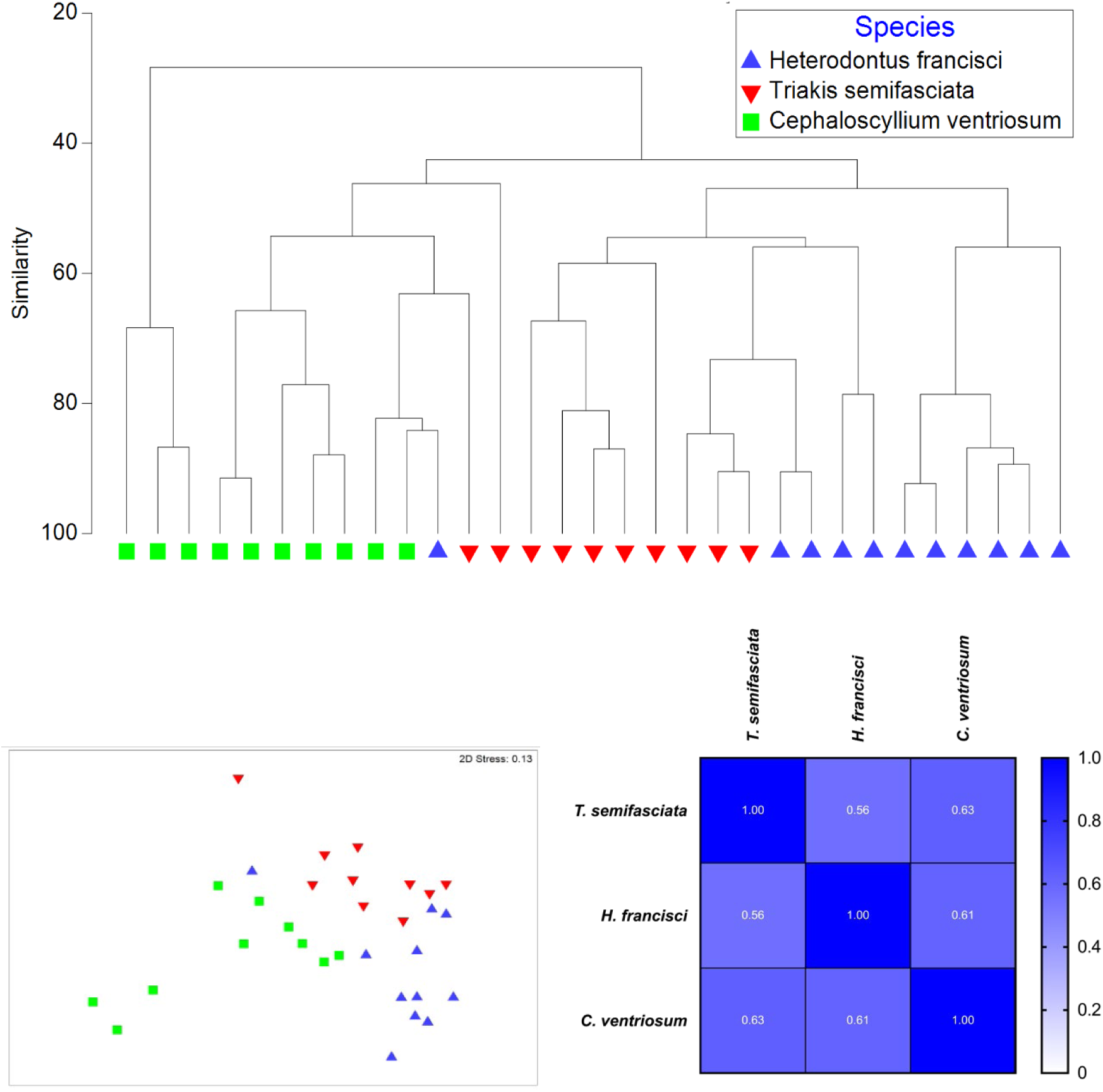
For *Heterodontus francisci*, *Triakis semifasciata*, and *Cephaloscyllium ventriosum*: A) Hierarchical clustering of epidermal metagenomes based on Bray-Curtis dissimilarities derived from fourth root transformed and standardized abundances at the genus level. B) Centroid nMDS ordination plots illustrating spatial differences among epidermal microbiomes. C) Spearman’s correlation matrix showcasing correlation coefficients among microbial genera.

### Dermal Denticle Morphology and Proportional Abundance Trends

The three shark species had variable dermal denticle morphology (Figure 5). *T. semifasciata* had overlapping dermal denticles, which were slightly elongated, *H. francisci* had square and crown shape dermal denticle that were evenly spaced, whereas *C. ventriosum* dermal denticles were highly elongated and widely, but unevenly spaced. Therefore, the most remarkable contrast in the morphology of denticles was the spacing between each scale: *T. semifasciata* had the greatest overlap (-197 ± 61.2 µm) while *H. francisci* had a greater average distance between each placoid (226 ± 15.1 µm) and *C. ventriosum* denticles were arranged with the greatest interdenticle distances (426 ± 41.4 µm; Figure 5).

**Figure 5.**
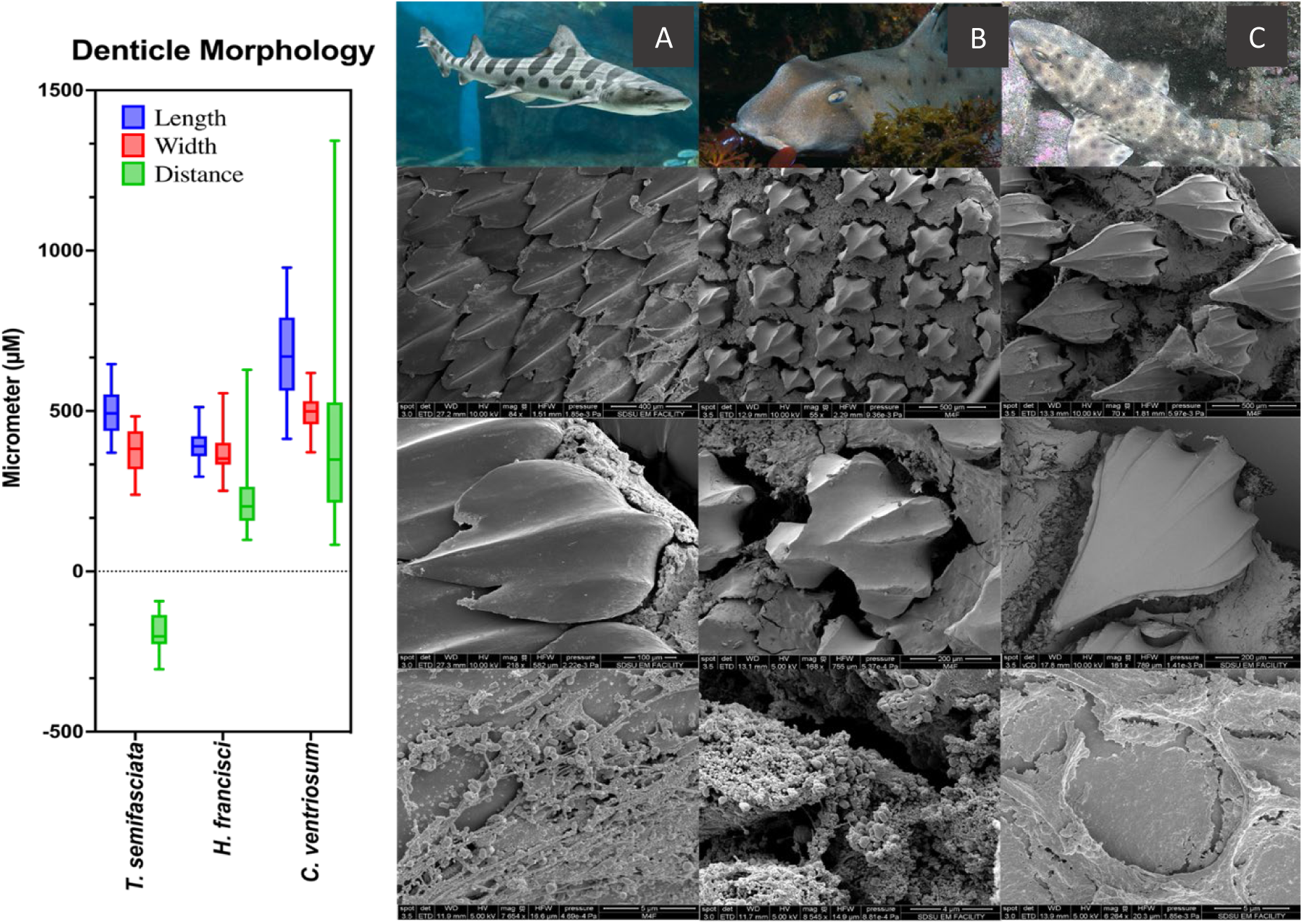
Box plot of denticle morphology depicting denticle length, width, and distance between denticles (left) measured and SEM images (right) at increasing magnification for A) *Triakis semifasciata*, B) *Heterodontus francisci* and C) *Cephaloscyllium ventriosum*. Magnification of SEM images in descending order is as follows: 500µm, 200µm, and 5µm.

In our investigation into the relationship between interdenticle distance and microbiome composition, we plotted the distances between denticles against the relative taxonomic abundances of the microbiome for each sampled shark species (Figure 6). Linear regression analysis revealed a moderately positive correlation for *Ehrlichia* (R² = 0.343, slope = 0.003) and *Portiera* (R² = 0.21, slope = 0.003). In contrast, *Bacteroides* exhibited a negative correlation (R² = 0.43, slope = -0.006) with increasing interdenticle distance. Both the positive and negative trends were statistically significant (*p* < 0.001), indicating weak to moderate linear relationships as evidenced by the correlation coefficients. No significant correlation (*p* > 0.05) was observed between interdenticle distance and the relative abundance of *Cyanothece* (R² = 0.031, slope = -0.011) when described by simple linear regression. Moreover, the relationship between denticle distance and both *Coprococcus* (R² = 0.324) and *Pseudoalteromonas* (R² = 0.44) were better described by second order polynomial equations.

**Figure 6.**
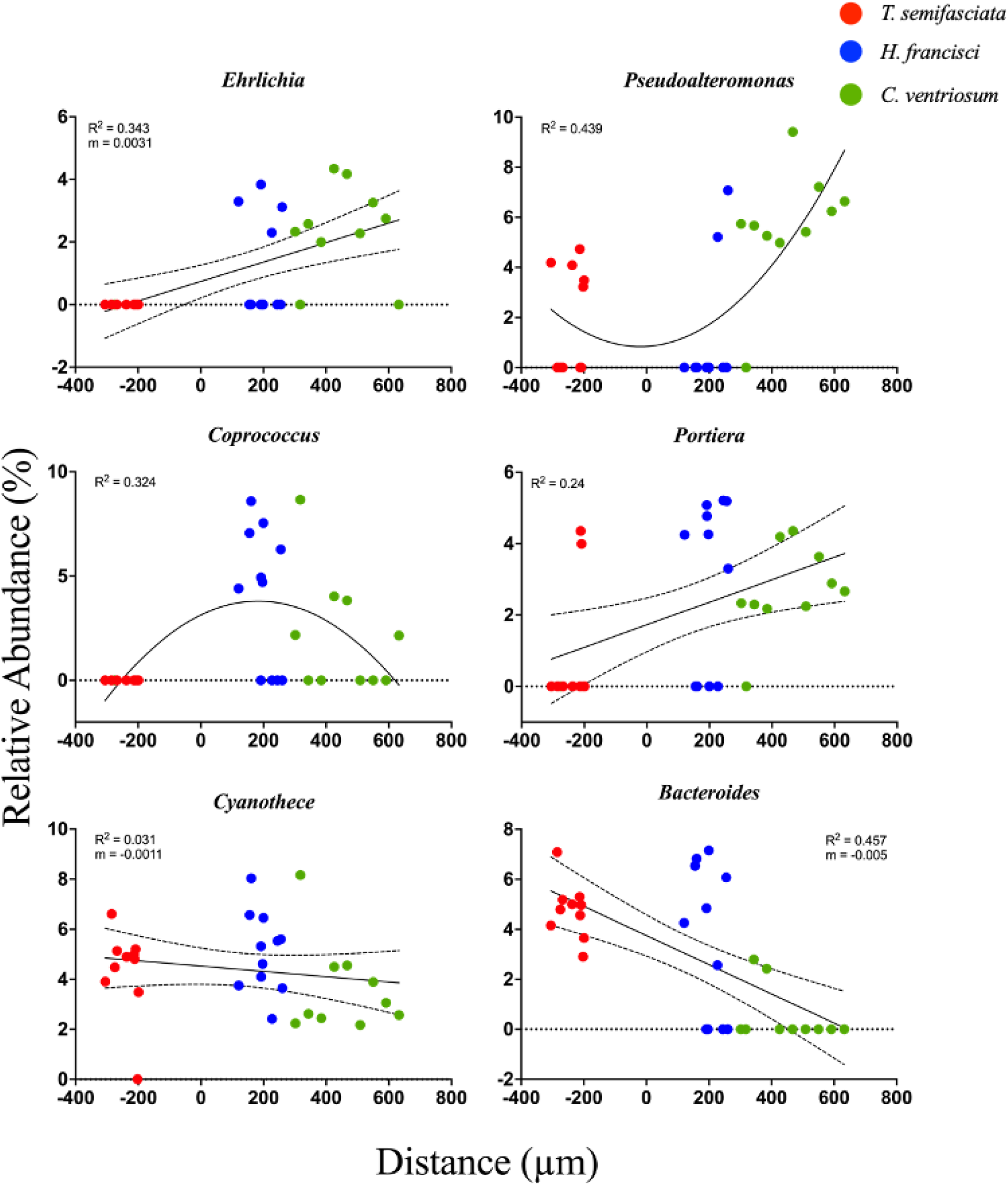
Scatter plots illustrating the correlation between interdenticle distances (x-axis) and microbial relative taxonomic abundance at the genus level (y-axis) for each shark sample. The curves represent the best fit from second-order polynomial regression analyses, and the associated goodness of fit (R²) and slope (m) values are displayed. The surrounding shaded areas define the confidence intervals for the best-fit curves.

### Functional Profiles of Epidermal Microbiomes Across Shark Species

Functional profiles of water-associated microbiomes collected from the captive environment were statistically dissimilar to host microbiomes in PERMANOVA main group tests (PERMANOVA: Genus, Pseudo-F _df= 1, 31_ = 3.21, P(perm) = 0.05) and pairwise analyses (*p* < 0.05) and are not discussed further. Of the 35 broadest functional genes, pathways involved in carbohydrate metabolism were most abundant (4.83 ± 0.1% S.D.), followed by amino acid synthesis (4.71 ± 0.1%), and protein metabolism (4.5 ± 0.26%). Although no significant difference overall was found between each pair of shark species (q = 0.85, p > 0.5), utilizing multiple Mann-Whitney tests, we compared the distributions of various functional genes across the three shark species (Table 5). For a more specific comparison, a Kruskal-Wallis H test revealed significant differences across the three species at the more specific functional level II including protein secretion system type 2 (p = 0.047), active compounds in metazoan cell defense (p = 0.023), general stress response (p = 0.04), lysine biosynthesis (p < 0.001), once-carbon metabolism (p < 0.001), and regulation of virulence (p = 0.039; all p-values corrected).

**Table 5.** Pairwise comparison of mean rank for functional pathways between three shark species: *T. semifasciata* (TS), *C. ventriosum* (CV), and *H. francisci* (HF). Table illustrates the p-values, mean ranks, mean rank differences, Mann-Whitney *U* statistics, and q-values, corrected using the two-stage step-up Benjamini, Kriekger and Yekutieli FDR method.

| Functional Pathway | P-value | Mean Rank |  |  | Mean Rank Diff. | Mann-Whitney U | q-value |
| --- | --- | --- | --- | --- | --- | --- | --- |
|  |  | HF | TS | CV |  |  |  |
| Cell Division and Cell Cycle | 6.8E-05 | 6.364 |  | 16.1 | -9.736 | 4 | 6.87E-04 |
| Cell Wall and Capsule | 5.5E-04 | 6.818 |  | 15.6 | -8.782 | 9 | 1.85E-03 |
| DNA Metabolism | 7.9E-04 | 6.909 |  | 15.5 | -8.591 | 10 | 1.99E-03 |
| Dormancy and Sporulation | 2.1E-03 | 7.182 |  | 15.2 | -8.018 | 13 | 3.47E-03 |
| Fatty Acids, Lipids, and Isoprenoids | 2.1E-03 | 7.182 |  | 15.2 | -8.018 | 13 | 3.47E-03 |
| Iron acquisition and metabolism | 2.8E-03 | 7.27 |  | 15.1 | -7.827 | 14 | 3.98E-03 |
| Membrane Transport | 8.0E-03 | 7.64 |  | 14.7 | -7.064 | 18 | 9.45E-03 |
| Miscellaneous | 2.8E-03 | 14.7 |  | 6.9 | 7.827 | 14 | 3.98E-03 |
| Motility and Chemotaxis | 2.1E-03 | 14.8 |  | 6.8 | 8.018 | 13 | 3.47E-03 |
| Nucleosides and Nucleotides | 2.6E-04 | 15.34 |  | 6.2 | 9.164 | 7 | 1.03E-03 |
| Phosphorus Metabolism | 1.7E-04 | 15.5 |  | 6.1 | 9.355 | 6 | 8.59E-04 |
| Photosynthesis | 1.7E-04 | 15.5 |  | 6.1 | 9.355 | 6 | 8.59E-04 |
| Potassium metabolism | 4.0E-05 | 15.7 |  | 5.8 | 9.927 | 3 | 6.87E-04 |
| Protein Metabolism | 6.8E-05 | 6.36 | 16.1 |  | -9.736 | 4 | 9.62E-04 |
| Respiration | 3.8E-04 | 6.73 | 15.7 |  | -8.973 | 8 | 1.54E-03 |
| Secondary Metabolism | 2.6E-04 | 15.4 | 6.2 |  | 9.164 | 7 | 1.44E-03 |
| Stress Response | 3.8E-04 | 15.3 | 6.3 |  | 8.973 | 8 | 1.54E-03 |
| Sulfur Metabolism | 2.8E-03 | 7.3 | 15.1 |  | -7.827 | 14 | 8.68E-03 |
| Virulence, Disease and Defense | 2.8E-03 | 7.3 | 15.1 |  | -7.827 | 14 | 8.68E-03 |
| Central metabolism | 1.1E-05 |  | 5.5 | 15.5 | -10 | 0 | 3.72E-04 |

To investigate the differences between the functional profiles of the metagenomes, we performed a SIMPER analysis on the sequenced genes at subsystem level II. Despite our focus on characterizing distinctions, we observed a high similarity between each species (SIMPER analysis: 100 - dissimilarity index), indicating a lack of pronounced differences at this level of analysis. Once again, the *T. semifasciata* group was most similar to *C. ventriosum* (93), while *H. francisci* was more dissimilar to both *C. ventriosum* (86) and *T. semifasciata* (85.3).

The functional gene potential of the microbiomes was significantly distinct across the three species (PERMANOVA, pseudo-F *_df=2,15_* = 2.47, p < 0.001; Table 6) at the broadest metabolic level. The difference between the three species was also detected when analyzing variations of more specific including SEED subsystem level II pathways (pseudo-F *_df= 2,15_* = 2.11, p < 0.05) and SEED subsystem level 3 (pseudo-F *_df= 2,15_*= 2.32, p < 0.05) metabolic pathways. Pairwise PERMANOVA also tests revealed consistently significant variations across species at each functional level (Table 7). However, no singular functional genes at level II subsystems differed between *T. semifasciata* and *H. francisci*.

**Table 6.** Summary of PERMANOVA and PERMDISP main test results across benthic shark epidermal microbiome functional subsystem levels I, II, and III (999 permutations).

| Gene Function: Level I | PERMANOVA |  |  |  |  | PERMDISP |  |
| --- | --- | --- | --- | --- | --- | --- | --- |
|  | df | SS | MS | Pseu<br>do-*F* | *P*-(perm) | *F*-value | *P*-(perm) |
| Benthic Sharks | 2 | 14.4 | 4.82 | 5.75 | **0.001** | 6.56 | **0.039** |
| Res | 28 | 11.7 | 0.837 |  |  |  |  |
| Total | 30 | 26.2 |  |  |  |  |  |
| **Gene Function: Level II** |  |  |  |  |  |  |  |
| Benthic Sharks | 2 | 47.1 | 23.6 | 3.22 | **0.001** | 10.8 | **0.007** |
| Res | 28 | 102.5 | 7.32 |  |  |  |  |
| Total | 30 | 149.6 |  |  |  |  |  |
| **Gene Function: Level III** |  |  |  |  |  |  |  |
| Benthic Sharks | 2 | 329.6 | 109.9 | 2.65 | **0.005** | 18.3 | **0.004** |
| Res | 28 | 580.7 | 41.5 |  |  |  |  |
| Total | 30 | 910.3 |  |  |  |  |  |

df = degrees of freedom, SS = sum of squares, MS = mean sum of squares.

**Table 7.**
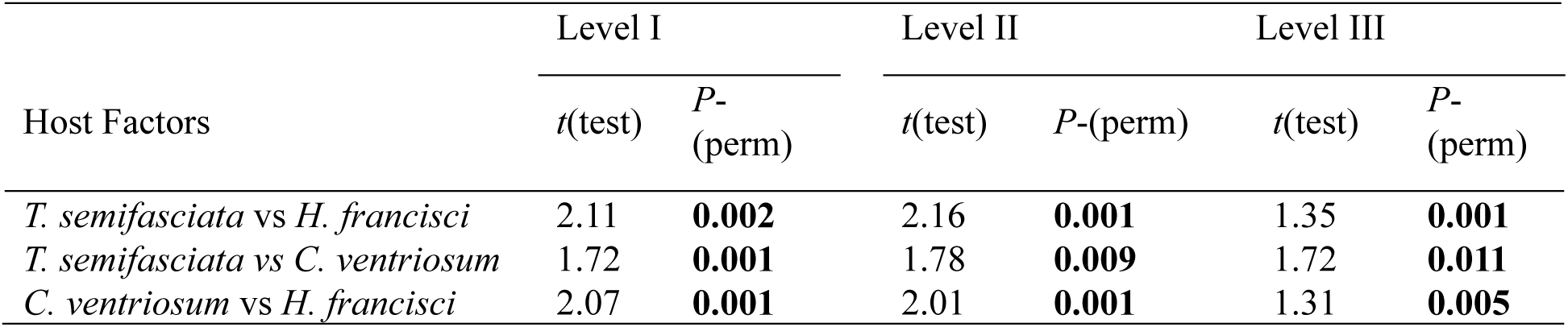
Summary of Pairwise PERMANOVA results between benthic shark species at functional subsystem levels I, II, and III (999 permutations).

| Host Factors | Level I |  | Level II |  | Level III |  |
| --- | --- | --- | --- | --- | --- | --- |
|  | *t*(test) | *P*-(perm) | *t*(test) | *P*-(perm) | *t*(test) | *P*-(perm) |
| *T. semifasciata* vs *H. francisci* | 2.11 | **0.002** | 2.16 | **0.001** | 1.35 | **0.001** |
| *T. semifasciata* vs *C. ventriosum* | 1.72 | **0.001** | 1.78 | **0.009** | 1.72 | **0.011** |
| *C. ventriosum* vs *H. francisci* | 2.07 | **0.001** | 2.01 | **0.001** | 1.31 | **0.005** |

In our exploration of the linear relationship between interdenticle distance and microbiome functionality, we plotted the distances between denticles against the relative abundance of Level II functional subsystems in the microbiome for each sampled shark species (Figure 7). Linear regression analyses outlined a significantly moderate, negative correlation (p < 0.001) between genes encoding for amino sugars (R^2^ = 0.38, slope = -0.004) and increase distance between denticles. The remainder of the significant relationships, notably those involving genes encoding chemotaxis (R^2^ = 0.18), regulation of virulence (R^2^ = 0.28), and desiccation stress (R^2^ = 0.54), electron-accepting reactions (R^2^ = 0.38), were more accurately characterized by second order (quadratic) curves, depicting a U-shaped distribution. Functional genes encoding for pyrimidine biosynthesis (R^2^ = 0.38) was characterized by a bell-shaped curve.

**Figure 7.**
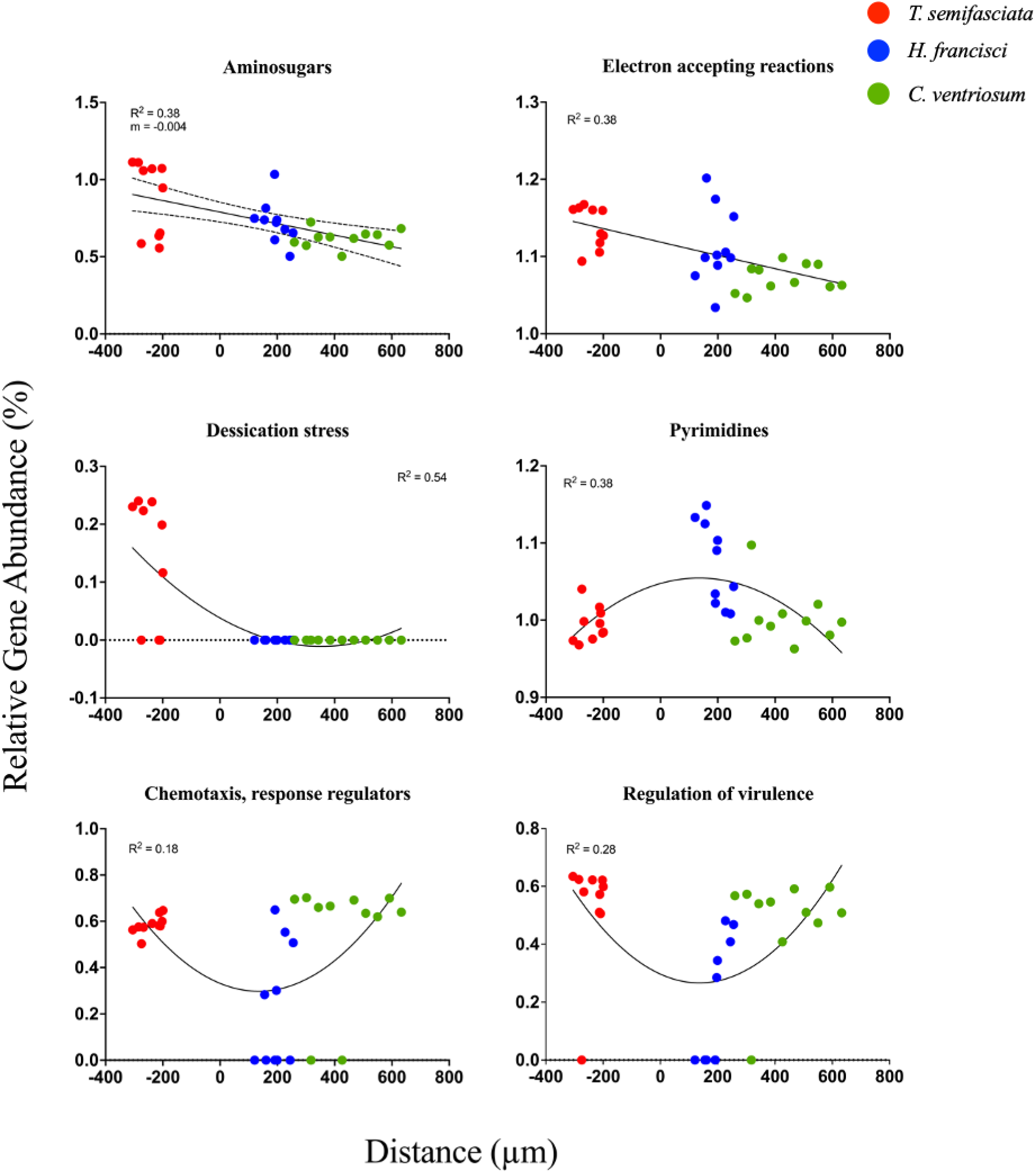
Scatter plots display the relationship between interdenticle distances (x-axis) and microbial relative functional gene abundance at SEED subsystem level 2 (y-axis) for each shark sample. The depicted best-fit lines are derived from both linear and second order polynomial regression analyses, each complete with their respective goodness of fit (R^2^) and slope (m).

The comprehensive multiple group comparisons did not uncover discernible linear correlations co-varying with the trajectory of denticle distance. However, these comparisons did provide evidence of host species-specific gene function levels. For example, the genes associated with desiccation stress were exclusively found in the microbiomes of *T. semifasciata*, with a significant average frequency (p < 0.05) of 0.113 ± 0.09 %. Notably elevated levels of genes associated with pyrimidine biosynthesis were discernible in the metagenomes of *C. ventriosum*, with a significant average frequency (p < 0.001) of 0.142 ± 0.1 % while a significant reduction (p < 0.01) in the levels of genes associated with virulence regulation in *H. francisci* microbiomes (0.17 ± 0.19 %) as compared to those observed in *T. semifasciata* (0.48 ± 0.2 %) and *C. ventriosum* (0.49 ± 0.1 %) was observed.

We investigated phylosymbiotic trends in both the taxonomic composition and functional gene profiles of the epidermal microbiomes associated with our three shark species. The theory of phylosymbiosis posits that as the evolutionary distance between host species increases — as determined by differences in the COX1 gene in our study — the dissimilarity of their associated microbiomes should also increase. This divergence can manifest in both the types of microbes (taxonomic dissimilarity) and in the functional genes those microbes carry (functional dissimilarity). Between the three species, *H. francisci* featured the farthest evolutionary distance from *C. ventriosum* and *T. semifasciata*, and we found significant increases in microbiome distance as the evolutionary distance increased (F *_df= 1,144_*= 5.1, Adj – R^2^ = 0.04, p < 0.05, Figure 8, left), supporting the argument for phylosymbiosis. The gene function comparisons corroborated the phylosymbiotic trends, with significant increase in dissimilarity again (F *_df= 1,144_* = 64.3, Adj-R2 = 0.3, p < 0.001, Figure 8, right).

**Figure 8.**
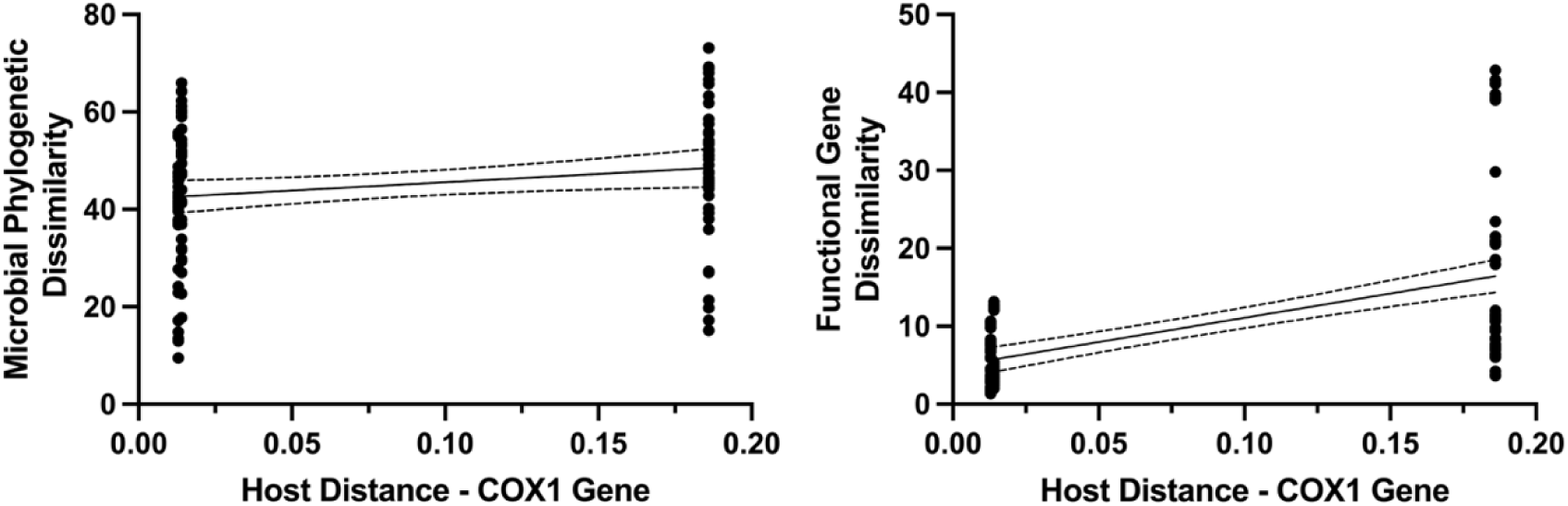
Phylosymbiotic comparison between host genetic divergence (COX1 gene) and epidermal microbiome phylogenetic dissimilarity (left) and gene function dissimilarity (right).

## Discussion

*Heterodontus francisci*, *Triakis semifasciata*, and *Cephaloscyllium ventriosum*, maintain an evolutionary distance of approximately 200 million years, with *T. semifasciata* and *C*. *ventriosum* belonging to the Carcharhiniform order and *H. francisci* of the Heterodontiformes and the host’s phylogenetic lineage shaped the epidermal microbiome, even when these benthic sharks inhabit the same coastal environment. Therefore, the unique morphological and physiological traits of the sharks directly impacted the relationship between host and epidermal microbiomes. Our research clarifies these intricate relationships, demonstrating that each shark species harbors a distinct microbial community characterized by both taxonomic composition and functional gene profiles. Interestingly, these microbiomes are influenced by the specific dermal denticle topography of the host, suggesting an intricate interplay between physical host characteristics and microbial colonization.

We evaluated the impact of host evolution across the sharks and observed a clear grouping of the *C. ventriosum* metagenomes in the plotted dendrogram, while *H. francisci* and *T. semifasciata* samples are more interspersed, suggesting more variability in the microbial communities of these species. However, consistent with the phylosymbiotic trend *T. semifasciata* and *C. ventriosum* metagenomes are more closely related to each other than *H. francisci* metagenomes. Moreover, metrics outlined by performing SIMPER analysis corroborate a significant co-evolution congruity in the epidermal microbiomes between *C. ventriosum* and *T. semifasciata* shark species. Finally, we found that as the evolutionary distance between host species increased, as measured by differences in the COX1 gene, the dissimilarity of their associated microbiomes also increased. This pattern was observed both in the taxonomic composition and functional gene profiles of the microbiomes, suggesting a deep intertwining of host evolution and microbiome development.

Building on phylosymbiotic trends, we found each shark species to harbor unique microbiome composition. For example, significant variation at phylum level, including differences in Pseudomonadota and Firmicute abundance between the three host species. While the skin of the shark species was dominated by Pseudomonadota, species-specific variations were observed among the Pseudomonadota classes Beta- and Gamma-proteobacteria. These distinctions echo a similar study finding fish, captive dolphins, whales, and killer whales which harbor species-specific microbiomes ^42^. Even though each host harbored a species-specific microbiome, we did observe some common features. Most notably, the Pseudoalteromonadaceae family was dominant across all shark hosts, consistent with microbiome surveys of *T. semifasciata* ^16^. This family of bacteria is well-known for its crucial roles in biofilm formation and the deterrence of potential microbial predator colonization^43,44^. An equally important contributor to the microbial compositions are the *Sphingomonodacaea* population which, belonging to the Alpha-proteobacteria clade, are key contributors to initial biofilm formation and are able utilize a wide array of organic compounds^45^. Together, these two major contributors dominate the microbiome of *T. semifasciata*, *H. francisci*, and *C. ventriosum*, and we theorize these phyla represent core contributors and regulators to the microbiome composition of elasmobranchs, being identified in multiple shark and ray species ^5,8,15,19^.

We investigated the impact of a host-derived factor, denticle topography, on the microbial compositions across the three species by quantifying the relationship between microbial abundance and interdenticle distance. We postulated that the overarching influence on microbial symbiotic associations across host species would be dictated by host phylogeny, with the effects of denticle topography exerting a selective influence on individual microbial constituents, contingent upon their unique bacterial characteristics, in a case-specific manner and found substantiating evidence for this theory. For example, the *Staphylococcus* genus was significantly more abundant in the metagenomes of *C. ventriosum*, less so in *T. semifasciata*, and completely absent in *H. francisci*. The associative pattern may be dictated by host phylogeny via differences in denticle coverage and subsequent mucus production across the three species. To better understand this phenomena we look to one study in which the attachment of two bacteria, *Escherichia coli* and *Staphylococcus aureus*, displayed varied attachment rates based on the surface of the synthetic models mimicking shark skin^46^. The attachment of *S. aureus* was inhibited on the surfaces resembling the highly structured shark denticle environment. Therefore, *Staphylococcus* associations may be selectively favored by the unique combination of decreased denticle coverage and increased mucus production characteristic of *C. ventriosum*. To build upon this theory, we posit the enhanced denticle overlap and less pronounced riblet elevation in conjunction with minimal but protein-rich mucus of *T. semifasciata* skin, provides a conducive surface for *Staphylococcus* to colonize^47^, while *C. ventriosum* offers the greatest opportunity for providing mucus production as a function of exposed epidermis. Last, the distinct structural environment of *H. francisci’s* dermal denticles, characterized by pronounced riblets and troughs, may be less hospitable to *Staphylococcus*, despite the potential assistance provided by mucus produced from a more exposed epidermal environment. Studies outline *Staphylococcus* species, such as *S. aureus*, to be considered common primary pathogens and implicated in wound infections^48–50^.

We understand coastal marine environments may serve as reservoirs of antibiotic-resistant strains of *Staphylococcus*, and the increased abundance across shark species may be indicators of increasingly opportunistic pathogenicity of associated microbiomes^51–53^. We observed similar phylosymbiotic trends between *Pseudoalteromonas* abundance and shark hosts, whereby host phylogeny served as a greater predictor of microbial association than respective denticle morphology. The associative behavior of the *Pseudoalteromonas* genus is a key contributor to biofilm regulation due to initial adhesion capabilities, subsequent pathogenic inhibition via the production of antibiofilm molecules^43,54,55^, and biosurfactants^56^. While the adhesion behavior of *Pseudoalteromonas* has not been investigated in the context of synthetic shark epidermal models, we can theorize this genus assumes the archetype of prolific biofilm initiator ^57^ and regulator ^58^ across all three species. Taken together, our observations underscore the intricate relationship between host denticle topography and microbial taxa, with *Staphylococcus* and *Pseudoalteromonas* serving as illustrative examples of how microbial colonization patterns are influenced by the host’s phylogenetic lineage via unique denticle topography.

Building on our observations, we were intrigued by the taxonomic abundances which echoed the denticle morphology. For example, *Ehrlichia*, often associated with pathogenic outcomes, and *Portiera*, known for its symbiotic relationships in insect hosts, both show positive covariance with increasing denticle distance. This association may reflect their adaptability to the physical environment provided by the denticle morphology, yet the specifics of this relationship are not well understood, and further research is warranted. It is therefore interesting to consider the ecological interaction between these bacteria and the composition of the shark denticles: denticle composition offers an ecological niche rich in organic matter including collagen fibers, which provide the scaffolding for the primary dentin mineral Hydroxyapatite^59^. Consequently, the moderate, negative covariance between *Bacteroides* and increasing interdenticle distance proved interesting given the wide distribution of the genus in marine degrading complex biopolymers, such as polysaccharides and proteins, which are integral to biofilm formation and organic carbon cycling^60^. The observed negative correlation between this genus and increasing interdenticle distance suggests that the decrease in denticle density may lead to a reduction in the available organic matter derived from enamel and dentine, substances that *Bacteroides* utilize, and therefore deter *Bacteroides* biofilm formation. Overall, the interplay between denticle morphology and bacterial associations is a complex, multifaceted relationship that extends beyond the confines of physical structure to include factors such as host physiology, including mucus production, and other environmental conditions.

The functional profiles of the epidermal metagenomes showed *T. semifasciata* microbiomes were most similar to *C. ventriosum*, while *H. francisci* was more dissimilar to both *C. ventriosum* and *T. semifasciata*, indicating functional redundancy and co-evolutionary trends and mirroring of phylogenetic distances between the species. We observed a significant, negative, covariance between genes encoding for both amino sugars and electron-accepting reactions with greater interdenticle distance. The observed relationship suggests that as denticle distance increases, microbes could be adapting their metabolic processes, specifically those related to amino sugar utilization, in response to the physical structure of the epidermis. This adaptation, reminiscent of how marine bacteria modulate their amino sugar production based on environmental cues, could involve an upsurge in the production of glucosamine and galactosamine, two pivotal amino sugars in bacterial physiology prevalent in marine ecosystems^61^. These linear trends establish a direct connection between shark skin morphology and specific functions of the epidermal microbiome. However, we also observed complex interplay between denticle distance and functional profiles of epidermal microbiomes when we mapped curvilinear relationships mirroring host phylogeny more closely than denticle morphology. For example, we observed U-shaped curves for chemotaxis, desiccation stress, and regulation of virulence, and bell curve line fitting pyrimidine biosynthesis, indicating a higher concentration of these genes at both the lower and higher extremes of interdenticle distance. This pattern of gene abundance, shaped by denticle distance, has notable implications for the functional potential of the microbiome, as these genes play critical roles in key microbial processes. Virulence regulation is implicated in disease-causing capabilities in bacteria in response to environmental cues including nutrient concentrations, pH, temperature, and host-derived factors^62^. The diminished representation of virulence regulation-associated genes within the epidermal microbiome associated with *H. francisci* implies heightened microbiome stability; a decreased necessity for bacterial pathogenesis could be indicative of a host-derived microbial community rather than an environmentally driven one, given habitats are conserved across the three species^63,64^.

Further investigation into each bacterial community’s tendency to follow or deviate from phylosymbiotic trends yielded insights into potential ecological interactions. For instance, genes related to desiccation stress were uniquely present in the microbiomes of leopard sharks, potentially due to the placement of goblet cells beneath the epidermis, resulting in a reduced mucus layer compared to other taxa. This suggests a heightened reliance on the dermal denticles for protection against desiccation. By highlighting functional gene differences, we further the knowledge about the influence of denticle distance for each shark host on associated bacterial populations. For example, the high denticle overlap covaried with elevated relative levels of genes encoding chemotaxis and flagellar movement suggests motile bacteria utilize the consistent and expansive area on or underneath the overlapping denticles instead of relying on the fluidics of the aqueous environment. The increased motility facilitated by flagella in the metagenomes can be theorized to allow bacteria to infiltrate and navigate biofilms, leading to a greater overall area of adhesion^65^. The relationship between bacterial motility and denticle overlap is invites varying interpretations. Although bacterial motility is not integral to biofilm initiation^66^, consistent surfaces presented by high overlap may facilitate biofilm formation, which requires swimming and twitching motility to navigate. Conversely, as denticle spacing increases, the necessity for enhanced bacterial motility could arise to navigate the intricate, aqueous environment, rather than relying on passive, random movement^65,67^. Further research is required to clarify the discriminating forces and reconcile these perspectives.

Previous efforts to understand these forces have already yielded insightful results. For instance, research conducted by Doane *et al*., which investigated the microbiomes of elasmobranchs of *Rhincodon typus*, *T. semifasciata*, and *A. vulpinus*, found no phylosymbiotic trends were observed within their functions ^19^. However, we detected phylosymbiotic patterns between the benthic species in this study both for microbial compositions and functional profile similarity. While we cannot completely dismiss the role of denticle distance or mucosal production as selective mechanisms, we posit the measured phylogenetic distance between *H. francisci* and both *T. semifasciata* and *C. ventriosum* is greater than that between *R. typus, T. semifasciata* and *A. vulpinus* and therefore, the pressure exerted by evolutionary distance is more evident. This theory is strengthened when considering sampling location as a driver of microbiome composition. If the environment exerted a greater pressure, we would expect to observe higher similarity between the benthic shark, and yet our findings revealed significant dissimilarities both between and within the groups.

## Caveats

Coupling core microbiomes of healthy animals with host-derived factors of microbiota recruitment will aid in the development of reliable biomarkers of shark ecology. However, although we have identified correlations between denticle topography and microbiome composition, our study design does not allow us to establish causality. Also, due to the high complexity of abiotic factors (e.g., water temperature, pH, salinity, and dissolved oxygen), and future threats of increased ocean temperatures, further research is needed to test the influence of environmental variables on the ability of marine hosts to recruit and retain epidermal microbes under extreme temperature shifts in controlled settings.

## Conclusion

In an exploration of the intricate relationship between host physiological characteristics and microbial community structures, we hypothesized the unique dermal denticles morphology of each shark species would parallel the influence of host phylogeny on the composition and structure of their corresponding epidermal microbiomes. Preliminary observations of epidermal microbiome taxonomic compositions and functional potentials differed between *T. semifasciata*, *H. francisci*, and *C. ventriosum* irrespective of the hosts sharing a captive environment. However, while our results reveal a compelling concordance between the diversity of epidermal microbiomes and host phylogeny among the examined shark species, we also observed consistent, yet weak, linear relationships with respect to denticle distance, a trait that does not follow the same phylogenetic pattern. This suggests a potential complex interplay between host evolutionary history and specific morphological traits in shaping the shark epidermal microbiome. The results of this study support the notion of phylosymbiosis, while suggesting denticle morphology provides a template for the assembly of microbial communities on shark skin.

## Materials and Methods

Epidermal microbiomes of captive *T. semifasciata* (*n* = 4), *H. francisci* (*n* = 3), and *C. ventriosum* (*n* = 10) were sampled in the summer of 2018 at the Birch Aquarium at Scripps Institution of Oceanography in La Jolla, California. In the summer of 2019, captive *T. semifasciata* (*n* = 6) were again sampled at the Birch Aquarium. Finally, in the summer of 2020, the epidermal microbiome of *H. francisci* (*n* = 8) were sampled in captivity at the National Oceanic and Atmospheric Administration (NOAA) in La Jolla, California. For all sampling events, captive sharks were immobilized in a sling for consistent collection of epidermal microbiomes located between the pectoral and dorsal fins above the lateral line on the left flank of each shark. Epidermal microbiomes were collected using a blunt, closed-circuit syringe prefilled with 100 kDa filtered seawater to flush the epidermis and displace microbes ^19,25,26^. Approximately 200 mL of captured microbes were then collected on a 0.22 µm sterivex, with one sterivex per individual. Water-associated microbial communities were collected using bulk water samples where approximately 60 L of tank water were simultaneously collected and first filtered through a nylon mesh sieve (200 µm pore size) to remove unwanted debris and eukaryotic organisms and second, concentrated using tangential flow filtration (100 kDa; ^27,28^ to produce ∼500 mL of tank water. The resulting concentration of tank water was filtered using a 0.22 µm sterivex.

Microbial cells anchored in sterivex filters were lysed by incubating the filters at 37 °C and 25 µL of proteinase K/SDS solution and resulting free DNA was extracted and purified using the Macherey-Nagel NucleoSpin Tissue Kit. The eluted DNA was prepared for shotgun metagenomic library sequencing using the Swift 2S Plus Kit (Swift Biosciences) and sequenced using an Illumina MiSeq sequencer. Samples were run in tandem using DNA barcoding throughout several sequencing runs as performed in previous studies ^26,27,29^. Resulting reads were processed for quality to remove artificial duplicates: reads with greater than 10 unknown nucleotides (n), and reads fewer than 60 base pairs (bp) in length via Prinseq++ ^30^. High quality, paired end reads were annotated via the Metagenomic Rapid Annotations using Subsystems Technology (MG-RAST; Keegan, Glass, and Meyer 2016) online database. MG-RAST calls taxonomic and functional gene assignments using BLAST comparisons to the National Center for Biotechnology (NCBI) and SEED genome databases ^32^. Sequencing annotations were conducted using the following parameters: e-value >10^-5^, 70 % identity, and > 60 bp alignment length.

Skin punches of 6 mm diameter were obtained from the dorsal flank regions of *H. francisci*, *T. semifasciata* and *C. ventriosum* sharks as metagenomic samples were collected. Biopsy specimen included the top layer of denticles and underlying dermal layers. Samples were rapidly frozen in liquid nitrogen (LN_2_) upon collection and maintained at cryogenic temperatures until fixed. Then, a 2.0 % glutaraldehyde (C_5_H_8_O_2_; Electron Microscopy Sciences, Hatfield, PA cat # 16100) and 0.1 M cacodylate buffer (C_6_H_12_AsNO_2_) was prepared fresh and the frozen skin punches dropped into the room temperature fixative. Samples were stored in fix at 4°C until prepared for scanning electron microscopy (SEM) analysis.

For SEM, tissues pieces were washed three times with 0.1 M cacodylate buffer to remove residual fix. They were further post-fixed in 1.0 % osmium tetroxide (OsO_4_; Electron Microscopy Sciences; Hatfield, PA cat #19150) in 0.1 M cacodylate buffer for 60 minutes at room temperature. Samples were dehydrated through a standard ethanol series of 30 %, 50 %, 75 %, two times 95 % and two times 100 % for 15 minutes each. Tissues pieces were critical point dried using liquid carbon dioxide (CO_2_) using a Samdri 790 CPD (Rockville, MD, USA). After drying, shark skin pieces were mounted with denticles facing up on 12mm Al stubs and double sticky carbon tape. The stubs were coated with ∼6nm platinum before viewing on FEI Quanta 450 SEM (Quanta 450, Hillsboro, OR USA) at 5-10kV.

For each shark species, the dimensions of the dermal denticles were measured. Dermal denticles were imaged at 100x magnification with a minimum of five fields per shark. On each image, five random denticles were selected and the widest and longest point of the denticle was measured. In addition, the distance to the nearest denticle was measured to provide an estimate of the inter-denticle distance.

### 2.4. Statistical Analyses

To study the impact of host phylogeny on the microbiomes of three captive shark species, we used several statistical analyses used historically in microbiome research^27,33,34^; first, to address the potential influence of rare taxa on the results, we fourth root transformed the data, followed by standardization ^28^, which is preferred to rarefaction ^35,36^. To assess the alpha-diversity of the microbial communities, we used Margalef’s (*d*), Pielou’s (*J’*), and Inverse Simpson’s (1/λ) indices to measure richness, evenness, and diversity, respectively ^37–39^. We tested the beta-diversity of the microbiomes by first comparing the microbiomes of sharks to the water column and second comparing between shark species using permutation multivariate analysis of variance (PERMANOVA). The PERMANOVA ran with 999 random permutations per analysis. To identify the similarities and differences between the groups, we calculated similarity percentage breakdowns (SIMPER)^40^. Mean rank comparisons were performed using the multiple comparison Friedman test, and to control the false discovery rate, the two-stage step-up method of Benamini, Krieger and Yekutieli was used. We conducted both PERMANOVA and non-parametric Kruskal-Wallis H tests on the relative abundance of microbial taxa from the levels of order to genus to examine changes in the microbiome’s taxonomy belonging to each host species. These tests were chosen as both do not assume a normal distribution and are appropriate for comparing three or more unrelated groups. Following community-level analyses, post-hoc analyses were performed to interpret the pairwise differences between the three shark species. In cases where the Kruskal-Wallis H test identified significant differences, the Tukey-Kramer post hoc test was applied to conduct pairwise comparisons between the groups. This post hoc test adjusts for multiple comparisons, reducing the likelihood of Type I errors. Last, a Bonferroni correction was applied to further control the family-wise error rate, adjusting the significance level to account for the multiple hypotheses tested. An alpha of 0.05 was used as the significance level for statistical tests.

To visualize the associations between metagenomes belonging to different but closely related species we generated non-metric multidimensional scaling (nMDS) derived from Bray-Curtis matrices ^41^. We chose an nMDS for its rank-based method of representing complex and non-linear relationships between multiple variable data. The Bray-Curtis dissimilarity was used as it effectively measures differences in community composition, accounting for the presence, absence, and abundance of species. A PERMDISP analysis was used to test for differences in group dispersion or homogeneity of the multivariate variations.

We simultaneously analyzed the abundance of genes in the microbiome as a proxy for gene expression. To assess the functional potential of the metagenomes of captive *H. francisci*, *T. semifasciata* and *C. ventriosum* sharks, we again used a PERMANOVA analysis. We also conducted an ANOVA with a post hoc Tukey test to identify differences in metabolism, which were visualized using the Statistical Analysis of Metagenomic Profiles (STAMP; v2.1.3; https://beikolab.cs.dal.ca/software/STAMP) software. All statistical analyses were performed using Primer-e package 7 (v7.0.2; accessed on 28 January 2022; www.primer-e.com/permanova.html) with the PERMANOVA+ add on, STAMP, and GraphPad PRISM 9 (v9.1.2; https://www.graphpad.com), and R studio. All graphs were generated using GraphPad PRISM 9. The SEED’s Subsystem Annotation was used to categorize the functional pathways into a hierarchical structure, ranging from broad metabolic pathways (Subsystem Level 1) to increasingly specific gene functions (Subsystem Levels II & III). This allowed us to map key biochemical functions to their parent pathways.

The correlation between microbial family relative abundance and the distance between shark denticles was examined using a regression model. To assess the strength and significance of the correlation, a scatter plot of the two variables was generated for each genus, along with respective p-value. The least squares regression line was superimposed on the scatter plot and the R-squared value was calculated, indicating how well the line fit the data.

To test whether skin microbiome composition was linked with host phylogeny, we calculated host distance by aligning the cytochrome c oxidase I (COX1) gene of each species using Clustal Omega on the EMBL-EBI server using default parameters. COX1 genes were downloaded from NCBI and used because it represents the only host gene publicly available for host phylogenetic comparison. We determined the relationship of host distance to microbiome similarity using linear modeling (lm; R)^19^.

## Institutional Review Board Statement

Animal handling and ethics were reviewed at San Diego State University through IACUC under permit APF #14–05-011D, APF #17–11-010D, APF # 18–05-007D, approved May 24th, 2018. Sampling was conducted under state permit SCP #12847 and SCP #9893 from the California Department of Fish and Wildlife.

## Acknowledgments

The authors would like to thank the Birch Aquarium at Scripps for allowing us to conduct research on site alongside staff. Students in the SDSU undergraduate ecological metagenomic class provide the sequencing of the leopard shark metagenomes.

## Data availability

All data and sequences were deposited in the NCBI Sequence Read Archive database.

